# CASB: A concanavalin A-based sample barcoding strategy for single-cell sequencing

**DOI:** 10.1101/2020.10.15.340844

**Authors:** Liang Fang, Guipeng Li, Qionghua Zhu, Huanhuan Cui, Yunfei Li, Zhiyuan Sun, Weizheng Liang, Wencheng Wei, Yuhui Hu, Wei Chen

## Abstract

Sample multiplexing facilitates single cell sequencing by reducing costs, revealing subtle difference between similar samples, and identifying artifacts such as cell doublets. However, universal and cost-effective strategies are rather limited. Here, we reported a Concanavalin A-based Sample Barcoding strategy (CASB), which could be followed by both single-cell mRNA and ATAC (assay for transposase accessible chromatin) sequencing techniques. The method involves minimal sample processing, thereby preserving intact transcriptomic or epigenomic patterns. We demonstrated its high labeling efficiency, high accuracy in assigning cells/nuclei to samples regardless of cell type and genetic background, as well as high sensitivity in detecting doublets by two applications: 1) CASB followed by scRNA-seq to track the transcriptomic dynamics of a cancer cell line perturbed by multiple drugs, which revealed compound-specific heterogeneous response; 2) CASB together with both snATAC-seq and scRNA-seq to illustrate the IFN-γ-mediated dynamic changes on epigenome and transcriptome profile, which identified the transcription factor underlying heterogeneous IFN-γ response.

## Introduction

Single-cell mRNA sequencing (scRNA-seq) and single-nucleus assay for transposase-accessible chromatin using sequencing (snATAC-seq) have emerged as powerful technologies for interrogating the heterogeneous transcriptional profiles and chromatin landscapes of multicellular subjects (Buenrostro, Wu et al., 2015, Cusanovich, Daza et al., 2015, Hashimshony, Wagner et al., 2012, Ramskold, Luo et al., 2012). Early scRNA/snATAC-seq workflows were limited to analyzing tens to hundreds of individual cells at a time. With the latest development of single-cell sequencing technologies based on microwells (Gierahn, Wadsworth et al., 2017), combinatorial indexing (Rosenberg, Roco et al., 2018) and droplet-microfluidics (Klein, Mazutis et al., 2015, Macosko, Basu et al., 2015), the parallel analysis of thousands of single cells or nuclei has become routine. The increase in throughput does not only lower the reagent costs per cell, but also enable the analysis of whole organs or entire organisms in one experimental run.

Recently, with the ever-increasing throughput, these technologies have also been used to reveal the temporal response of heterogeneous cell population under diverse perturbations, which require tens of samples to be processed in parallel (Hurley, Ding et al., 2020, Weinreb, Rodriguez-Fraticelli et al., 2020). Based on existing methods, sample-specific barcodes (for example, Illumina library indices) are often incorporated at the very end of standard library preparation workflow. Such workflow requires parallel processing of multiple individual samples until the final step, therefore not only is labor-intensive and limits the number of samples, but also increase the reagent costs if a small number of cells would be sufficient to characterize the heterogeneity of each individual sample. To overcome this, alternative multiplexing approaches should label cells from each sample with distinct barcodes before pooling for single-cell sequencing experiment. The sample-specific barcodes could then be linked to cell barcodes during single-cell sequencing library preparation. Several methods have been developed in this endeavor, which introduce sample barcodes using either genetic or non-genetic mechanisms. Genetically, researchers have used various strategies to express an exogenous gene with sample-specific barcodes at its 3′ UTR, which can be captured similarly as endogenous genes (Hurley et al., 2020, Weinreb et al., 2020); non-genetically, people have used oligonucleotide containing a sample barcode followed by a poly-A sequences, which can be immobilized on the cell or nuclear membrane through anchoring molecules (e.g. antibody and lipid) (McGinnis, Patterson et al., 2019, Stoeckius, Hafemeister et al., 2017) or chemical cross-linking reaction (Gehring, Hwee Park et al., 2020), or defused into permeabilized nuclei (Srivatsan, McFaline-Figueroa et al., 2020), and then captured during reverse transcription. Although being already quite powerful, each of these methods has still its own liabilities, including issues with scalability, universality or the potential to introduce artefactual perturbations. Moreover, all of these methods have only been combined with scRNA/snRNA-seq, and have not yet been applied and are likely incompatible with snATAC-seq.

Here, we developed a Concanavalin A-based Sample Barcoding strategy (CASB) that overcomes many of these limitations. Taking advantage of the glycoprotein-binding ability of concanavalin A (ConA), CASB was used to label cell or nucleus with biotinylated single-strand DNA (ssDNA) through a streptavidin bridge. CASB could be easily adapted into scRNA/snATAC-seq workflows, and showed high accuracy in assigning cells or nuclei regardless of genetic background as well as in resolving cell doublets. The application of CASB in samples with time-series experiments, followed by scRNA- and/or snATAC-seq, allows revealing diverse transcriptome/epigenome dynamics.

## Results

### CASB enables cell and nucleus labeling with ssDNA

The CASB consists of three components: biotinylated ConA, streptavidin and biotinylated ssDNA as barcoding molecules. Both ConA and streptavidin form homo-tetramer autonomously, allowing the assembly of ConA-streptavidin-ssDNA complex (Fig 1A). Relying on the glycoprotein-binding ability of ConA, such assembled complex can be immobilized on the cell or nuclear membrane (Fig 1A). To measure how many ssDNA molecules can be immobilized on the cell membrane, a biotinylated ssDNA with 5′ and 3′ PCR handles flanking an eight-nucleotide (N8) random sequence was used to label the cells (Fig 1A). After incubation with different quantity of preassembled ConA-streptavidin-ssDNA complex in PBS at 4 °C (Methods), the number of ssDNA molecules immobilized on mouse embryonic stem cells (mESC) was quantified using qPCR. As shown in Figure 1B, the amount of ssDNA immobilized on cells increased with the increased usage of ConA-streptavidin-ssDNA complex, and could reach as many as 50,000 molecules per cell. To test whether ssDNA may fall off from labeled cells and cause cross-contamination during sample pooling, a mouse embryonic fibroblast (MEF) cell population expressing mCherry fluorescent proteins was labeled with the ssDNA and then mixed with another MEF cell population expressing GFP fluorescent proteins, which was only coated with “empty” ConA (Methods). After 30 min incubation in PBS at 4 °C, mCherry and GFP positive cells were separated using FACS and subjected to qPCR measurement. As shown in Figure 1C, the ssDNA immobilized on mCherry+ cells was not detectable from GFP+ cells, demonstrating the stability of CASB labeling. In addition to labeling the whole cell, we also measured the labeling efficiency of CASB for cell nucleus, in which nuclei were labeled with preassembled ConA-streptavidin-ssDNA complex in nuclear extraction buffer at 4 °C (Methods). As shown in Figure EV1A, the amount of ssDNA immobilized on nuclei increased with the increased usage of ConA-streptavidin-ssDNA complex, and reached at least 120,000 molecules per nucleus. Taken together, these results demonstrated that CASB is able to stably label both cell and nucleus with ssDNA.

**Figure 1.**
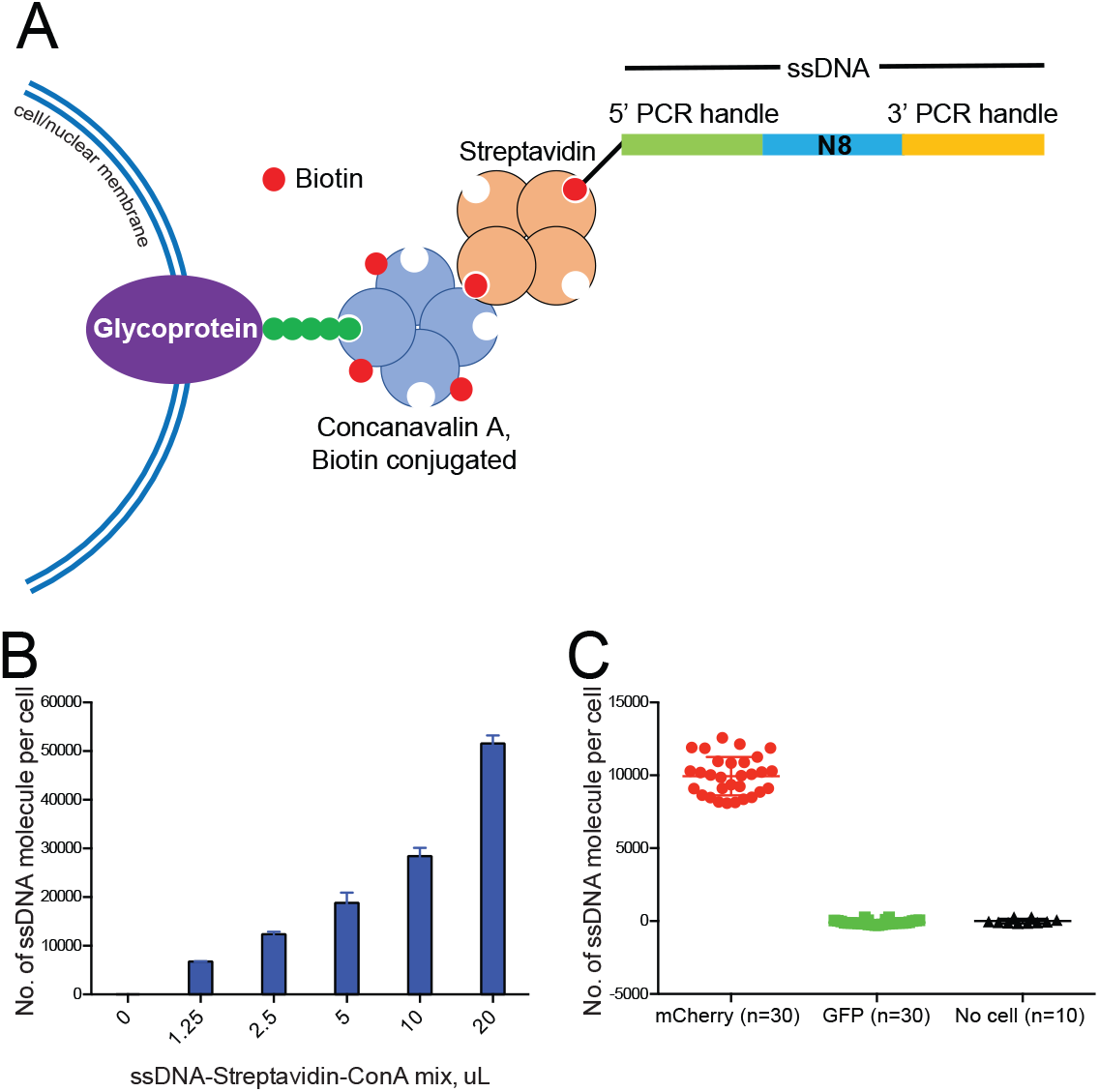
Cell labeling with CASB. **A**, An illustration of CASB. Biotinylated ssDNA was immobilized on glycoprotein on cell/nuclear membrane through streptavidin and biotinylated ConA. The ssDNA contains 5′ and 3′ PCR handles that flank an 8 nt random sequence. **B**,mESC were labeled with different quantity of CASB, and the number of ssDNA molecules immobilized on mESC was quantified used qPCR. The amount of ssDNA immobilized on cells increased with the increased usage of ConA-streptavidin-ssDNA complex, and reach as many as 50,000 molecules per cell. **C**,CASB-labeled mCherry+ MEF cells were incubated with unlabeled GFP+ MEF cells. The number of ssDNA molecules immobilized on mCherry+ and GFP+ cells was quantified used qPCR after FACS separation. The ssDNA immobilized on mCherry+ cells was not detectable from GFP+ cells. “n” means number of qPCR reactions. Error bars represent SD.

### CASB enables scRNA-seq sample multiplexing

In scRNA-seq, cell specific barcodes are attached to the cDNA during reverse transcription (RT) by using primers consisting of a cell barcode sequence, a unique molecular identifier (UMI) sequence and a poly-T sequence that anchors to the poly-A tail of mRNA molecule. To make our CASB compatible with the standard scRNA-seq workflow, we designed a biotinylated barcoding ssDNA with a 5′ PCR handle followed by a N8 barcode and a 30 nt poly-A tail, which can be captured by a RT primer consisting of a PCR handle followed by a 30 nt poly-T tail (Fig 2A). After CASB labeling, MEF cells were directly lysed and subjected to RT reaction (Methods). The barcoding ssDNA immobilized on cell membrane was quantified together with the endogenous housekeeping gene ActB using qPCR. As shown in Figure EV1B, both barcoding ssDNA and ActB gene can be efficiently capture by RT primer. Therefore, CASB could be easily adapted into scRNA-seq workflow with high efficiency.

**Figure 2.**
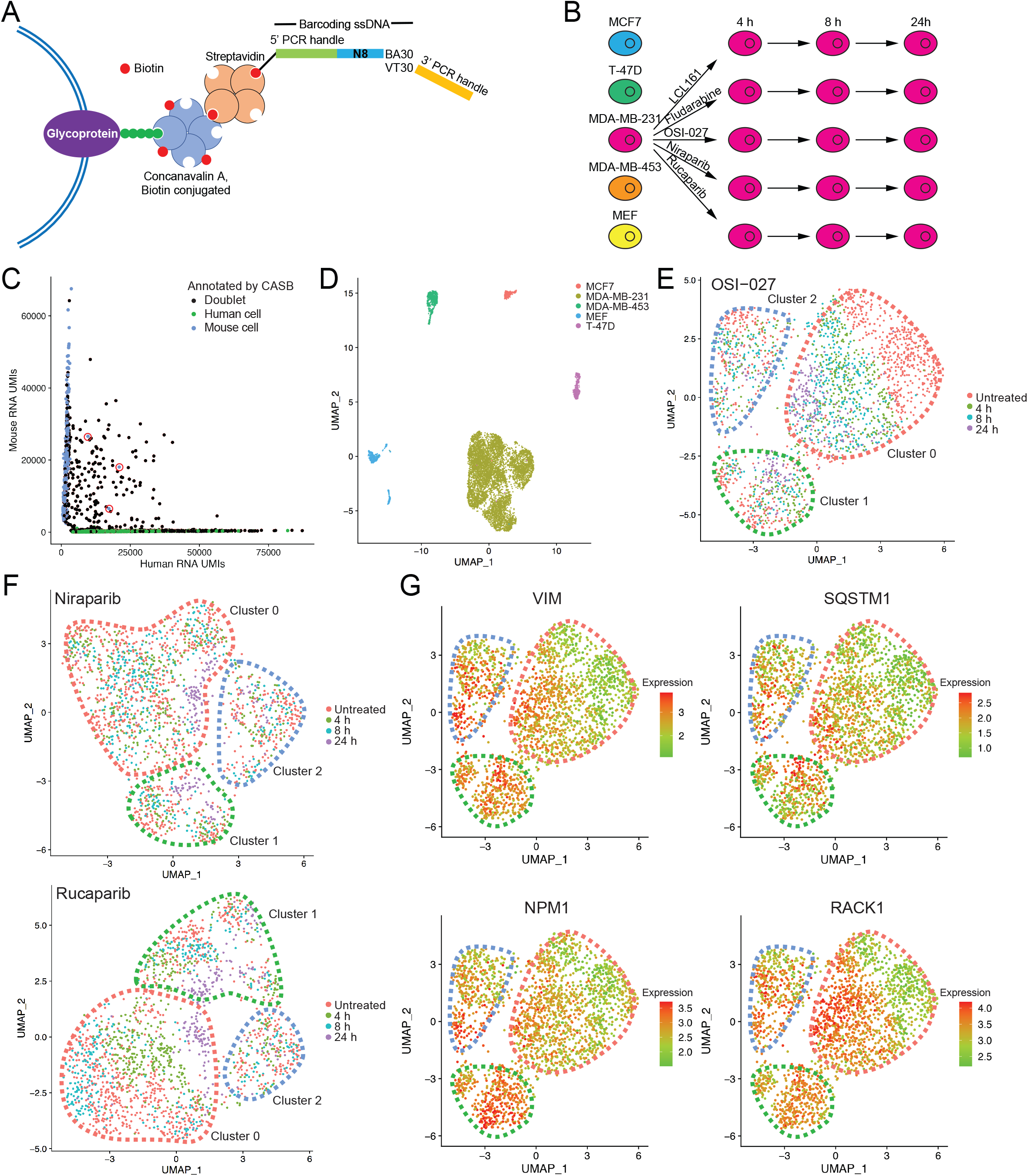
CASB enables scRNA-seq sample multiplexing. **A**, An illustration of CASB used in scRNA-seq. A biotinylated barcoding ssDNA with a 5′ PCR handle followed by an 8 nt barcode and a 30 nt polyA tail was used to mimic the endogenous transcripts. **B**,The design of the experiment. MDA-MB-231 cells were perturbed with 5 different compounds, collected at 3 different time points, CASB labeled, and then pooled with 3 other breast cancer cell lines and MEF cells. **C**,Scatter plot depicting the number of UMIs associated with transcripts from human or mouse genome. Cell doublets revealed by CASB were marked in black. Out of 110 mouse-human doublets, 107 were detected as doublets by CASB barcodes. Three interspecies cell doublets that were not detected by CASB were circled in red. Beside interspecies cell doublets, cell doublets from one species were also detected by CASB. **D**,Transcriptome-based UMAP of cells captured in scRNA-seq. Cells were colored according to the CASB barcodes, and doublets were excluded. Different human and mouse cells formed 5 distinct cell clusters, respectively. **E&F**,Transcriptome-based UMAP of untreated and (**E**) OSI-027-, (**F**) Niraparib- and Rucaparib-treated MDA-MB-231 cells. Three cell populations with distinct transcriptomic responses were observed in each UMAP: sensitive cell subpopulation was circled in red, while insensitive ones in green and blue, respectively. **G**,Transcriptome-based UMAP of untreated and OSI-027-treated MDA-MB-231 cells. Sensitive cell subpopulation was circled in red, while insensitive ones in green and blue, respectively. Expression level of VIM, SQSTM1, NPM1 and RACK1 are indicated by color code, which were expressed in untreated insensitive cell populations and induced by OSI-027 in sensitive cells.

To demonstrate the strength of CASB in scRNA-seq, a breast cancer cell line MDA-MB-231 was perturbed with 5 different compounds, collected at 3 different time points after treatment and pooled with 3 other breast cancer cell lines as well as MEF cells after separate sample labelling using CASB (Fig 2B). Unlabeled MDA-MB-231 cells were also added into the sample pool to measure the potential influence of CASB on transcriptome profile. Sample pool was then subjected to scRNA-seq using the 10x Genomics Chromium system with minor modifications: 1) in order to examine the efficiency of CASB to detect doublets, we intentionally overloaded the system (~20,000 instead of ~10,000 cells recommended by the manufacturer) to create more cell doublets; 2) CASB barcode and transcriptome library were separated by size selection before next-generation sequencing library construction, enabling pooled sequencing at user-defined proportions (Methods).

As a result, a total of 12068 cells were captured with sufficient reads for transcriptome analysis. For each cell, the reads derived from each of the 20 different sample barcodes were counted and used to demultiplex the samples using HTODemux method (Stoeckius, Zheng et al., 2018) (Methods). A total of 483 cells were assigned as ‘Unlabeled’, as expected due to the inclusion of unlabeled MDA-MB-231 cells (Fig EV1C). Among the remaining ones, 3962 cells were assigned as cell doublets encapsulated in the same droplet, as they contained two or three major barcodes (Fig EV1C). Indeed, the doublets consisting of both mouse and human cells, which could be unambiguously detected based on their mapping results, could also be efficiently identified based on the mixture of CASB barcodes. As shown in Figure 2C, out of 110 mouse-human doublets, 107 (97.3%) were defined as doublets based on our CASB data. When compared with singlets, more UMI derived from both CASB barcode and mRNA transcripts were detected in doublets (Fig EV1D), further validating the correct assignment of cell doublets. Within 7623 singlets, the number of detected UMI from CASB per cell ranged from 245 to 2134 (5-95 percentile), and significantly correlated with UMI detected for endogenous transcripts in the same cell (Fig EV1E), suggesting a similar cell-specific capture efficiency between CASB barcode and endogenous transcripts, and that CASB did not impair mRNA capture. Based on the qPCR quantification, the same amount of CASB mixture could label cells with ~20,000 ssDNA (Fig 1B), mean UMI (1051) detected in the scRNA-seq indicates a ~5% capture efficiency at current sequencing depth (25 million total sequencing reads). To determine the variation of labeling efficiency among different cells, given the cell-specific capture efficiency, we first normalized the number of UMI numbers from CASB by that of UMI derived from the endogenous transcripts in the same cell. As shown in Figure EV1F, the CASB barcoding manifested a good uniformity of labeling efficiency among all singlets and within individual cell samples. Taken together, our CASB strategy could achieve high sensitivity in detecting cell identity and doublets in scRNA-seq experiments.

For the 7623 cells with unambiguously assigned sample origin, we then clustered them based on their scRNA-seq profiles. As shown in Figure 2D, different human and mouse cells formed 5 distinct cell clusters, respectively. Each cluster was composed of cells from individual cell line labeled with distinct CASB indices (Fig 2D). We also compared the untreated MDA-MB-231 cells with to those without CASB labelling. As shown in Figure EV1G and H, single cell profiles were intermingled together and their cumulative transcriptome were highly correlated, demonstrating a negligible influence of CASB labeling on transcriptome profile. Within the MDA-MB-231 cell population, all 16 sample barcodes can be detected (Fig EV1I). Cells associated with 24 h-treatment of Niraparib, Rucaparib and OSI-027 could be well distinguished from untreated cells, whereas those with LCL161 and Fludarabine could not (Fig EV1J). As expected, Niraparib- and Rucaparib-treated cells were intermingled due to their common molecular target PARP.

MDA-MB-231 is of triple-negative breast cancer origin, which lacks efficient targeted therapy. As intratumoral heterogeneity has been associated with therapy resistance, we investigated whether drug treatments could lead to heterogeneous response in the MDA-MB-231 cells. We focused on compound OSI-027, as it induced the largest transcriptomic changes (Fig EV1J). As shown in Figure 2E and EV2A, in which cells treated with OSI-027 were plotted with untreated cells, there indeed existed three cell populations with distinct transcriptomic responses. Whereas one showed clearly time-dependent transcriptomic changes (Fig 2E, cluster 0 circled in pink), the other two had limited alteration in gene expression even after 24 h (Fig 2E, cluster 1&2 circled in green and blue, respectively), suggesting that the latter were less sensitive to the OSI-027. Neighbor proportion analysis also confirmed that, untreated cells were well separated from treated cells in cluster 0, while it is not the case for cluster 1 and 2 (Fig EV2B). As cellular heterogeneity existed already in the untreated MDA-MB-231 cells (Fig EV2C), we sought to further check whether the insensitive cell populations were also resistant to other effective compounds. Indeed, as shown in Figure 2F and EV2D, the same two cell clusters also appeared less sensitive to Niraparib and Rucaparib, suggesting the intrinsic multidrug insensitivity. To explore the underlying factors, we perform function enrichment analysis on genes that were commonly up- or downregulated in cluster 1 and 2 compared to cluster 0 using IPA software (Methods). Interestingly, these genes were highly enriched in the cellular compromise and movement pathways (Table EV1 & Fig EV2E). Importantly, many of them (e.g. VIM) were upregulated in cluster 1 and 2, and known to promote cellular movement (Fig EV2F). In tumor cells, increased cell motility mediated by epithelial-mesenchymal transition (EMT) is also highly associated with drug resistance (Singh & Settleman, 2010, Zhang & Weinberg, 2018). Our results suggested that the intrinsic multidrug insensitivity of MDA-MB-231 cells may result from the activated EMT. More interestingly, when overlap the potential insensitivity-causing genes in cluster 1 and 2 with OSI-27-regulated genes, we observe that many genes, including VIM, SQSTM1, NPM1 and RACK1, were also upregulated by OSI-027 in sensitive cells (Fig 2G). Given these genes are involved in promoting EMT and potentially also drug resistance (Fig EV2F), this observation indicated the potential of OSI-027 treatment in inducing acquired therapy resistance.

### CASB enables snATAC-seq sample multiplexing

So far, no sample multiplexing method has been developed for droplet-based snATAC-seq. In droplet-based snATAC-seq, cell nuclei are firstly incubated with transposase in bulk, where genomic DNA is fragmented and tagged with adapter sequences. Afterwards, single cells are encapsulated, and cell barcodes are added to DNA fragments during PCR in individual droplets using primers targeting the adapter sequences. To adapt CASB into snATAC-seq workflow, we designed a 222 nt barcoding ssDNA with S5-ME and S7-ME adapter sequences flanking a sequence containing sample barcodes (Fig 3A). S5-ME and S7-ME adapter sequences were used as primer anchoring sites during snATAC-seq library amplification (Methods). The labeling efficiency using such ssDNA was measured similarly as before, in which nuclei were directly labeled with preassembled ConA-streptavidin-ssDNA complex in nuclear extraction buffer at 4 °C (Methods). As shown in Figure EV3A, the amount of ssDNA immobilized on nuclei increased with the increased usage of ConA-streptavidin-ssDNA complex, and could reached at least 80,000 molecules per nucleus.

**Figure 3.**
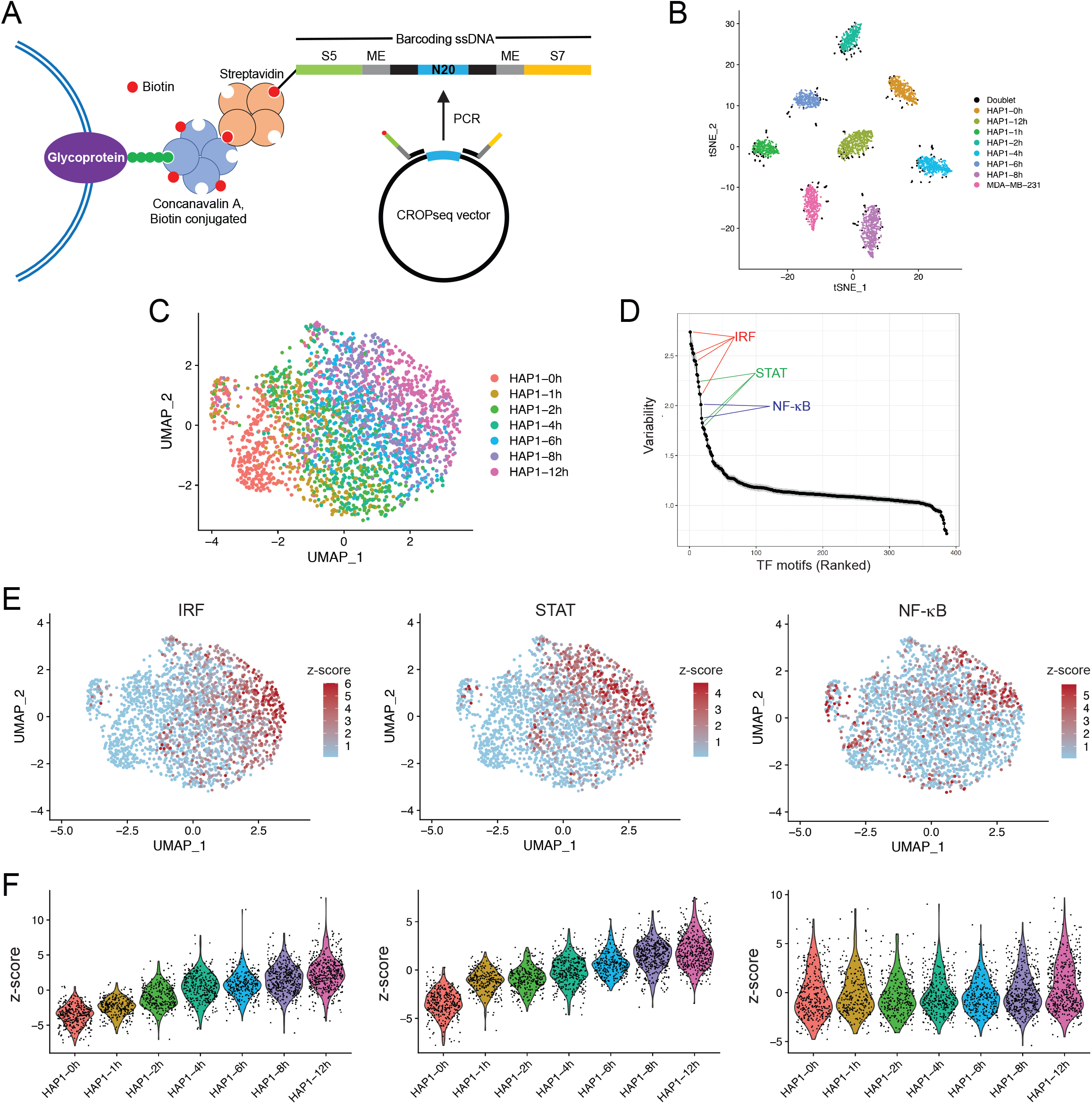
CASB enables snATAC-seq sample multiplexing. **A**, An illustration of CASB used in snATAC-seq. A biotinylated barcoding ssDNA with S5-ME and S7-ME adapter sequences flanking a sequence containing sample barcodes was used to mimic the transposed genomic DNA. **B**,t-SNE projection based on the CASB barcode reads captured in snATAC-seq. Cells were colored according to the CASB barcodes, and doublets were marked in black. **C**,ATAC-based UMAP of all HAP1 cells captured in snATAC-seq. Cells were colored according to the CASB barcodes, and doublets were excluded. HAP1 cells showed a continuous shift in chromatin profile from 0 to 12 h. **D**,Dot plot revealing the TFs with the most variable activity across all cells including IRF, STAT and NF-κB. **E**,ATAC-based UMAP of all HAP1 cells, in which the TF activity was presented by bias-corrected deviation z-score across all cells in color code. **F**,Violin plots demonstrating the deviation z-score of different TFs across different cells at different time points. IRF and STAT associated peaks showed continuous activation upon IFN-γ stimulation, while the variation of NF-κB peak intensity was largely due to the heterogeneity within HAP1 cells.

To demonstrate the application of CASB in snATAC-seq, we sought to monitor the temporal chromatin changes induced by interferon-gamma (IFN-γ) in HAP1 cells. IFN-γ is an important cytokine in the host defense against infection by viral and microbial pathogens (Shtrichman & Samuel, 2001). It mediates innate immunity through regulating effector gene expression, which is accompanied by substantial changes at epigenetic level (Ivashkiv, 2018). However, how heterogeneously and dynamically cells respond to IFN-γ stimulation at the chromatin level has remained elusive. Taking advantage of CASB, we analyzed the changes in chromatin accessibility of HAP1 cells at 7 different time points after IFN-γ stimulation using snATAC-seq. MDA-MB-231 cells were added into the pool here as an outlier control. After sequencing, a total of 2890 cells were obtained with sufficient reads, 305 of which were identified as cell doublets that have at least two major CASB barcodes (Fig 3B, marked in black), and 23 cells were unlabeled (Fig EV3B). MDA-MB-231 cells with its specific CASB barcode presented as an isolated cluster (Fig EV3C). As revealed by UMAP projection, HAP1 cells showed a continuous shift in chromatin profile from 0 to 12 h (Fig 3C). To uncover the key transcription factors (TFs) that mediate IFN-γ-induced chromatin remodeling, we analyzed their binding motifs on the ATAC-peaks across all cells and observed that peaks containing motifs of IRF, STAT and NF-κB TF showed large variation in their intensity, indicating their functions in modulating IFN-γ response (Fig 3D) (Methods). Indeed, IRF and STAT peaks showed continuous activation (Fig 3E&F). This is expected as IFN-γ is able to activate JAK/STAT signaling through binding to its receptor, which in term activates the expression of IFN-γ-responsive genes, including transcription factors IRFs (Leonard & O’Shea, 1998). Interestingly, the large variation of NF-κB peak intensity did not result from IFN-γ treatment, but was instead largely due to the heterogeneity of HAP1 cells (Fig 3F). It is known that NF-κB can be activated by IFN-γ and able to facilitate the transcription activation of IFN-γ targets, including CXCLs (Pfeffer, 2011, Qin, Roberts et al., 2007). This result suggested that heterogeneous NF-κB activity may give rise to heterogeneous IFN-γ response.

To evaluate whether heterogeneous NF-κB activity causes heterogeneous IFN-γ response, IFN-γ treated samples from the same time points were also analyzed using CASB followed by scRNA-seq. A total of 3407 cells were captured, 294 of which were identified as cell doublets and 9 cells were unlabeled (Fig EV4A&B). As shown in Figure 4A, HAP1 cells showed a continuous shift in the transcriptome profile from 0 to 12 h on UMAP projection. To globally evaluate the correlation between chromatin accessibility and gene expression, we analyzed the dynamic expression patterns of predicted IRF and STAT target genes. In consistent with the activity of IRF and STAT observed in snATAC-seq (Fig 3C), the expression of their target genes also exhibited continuous upregulation (Fig 4B).

**Figure 4.**
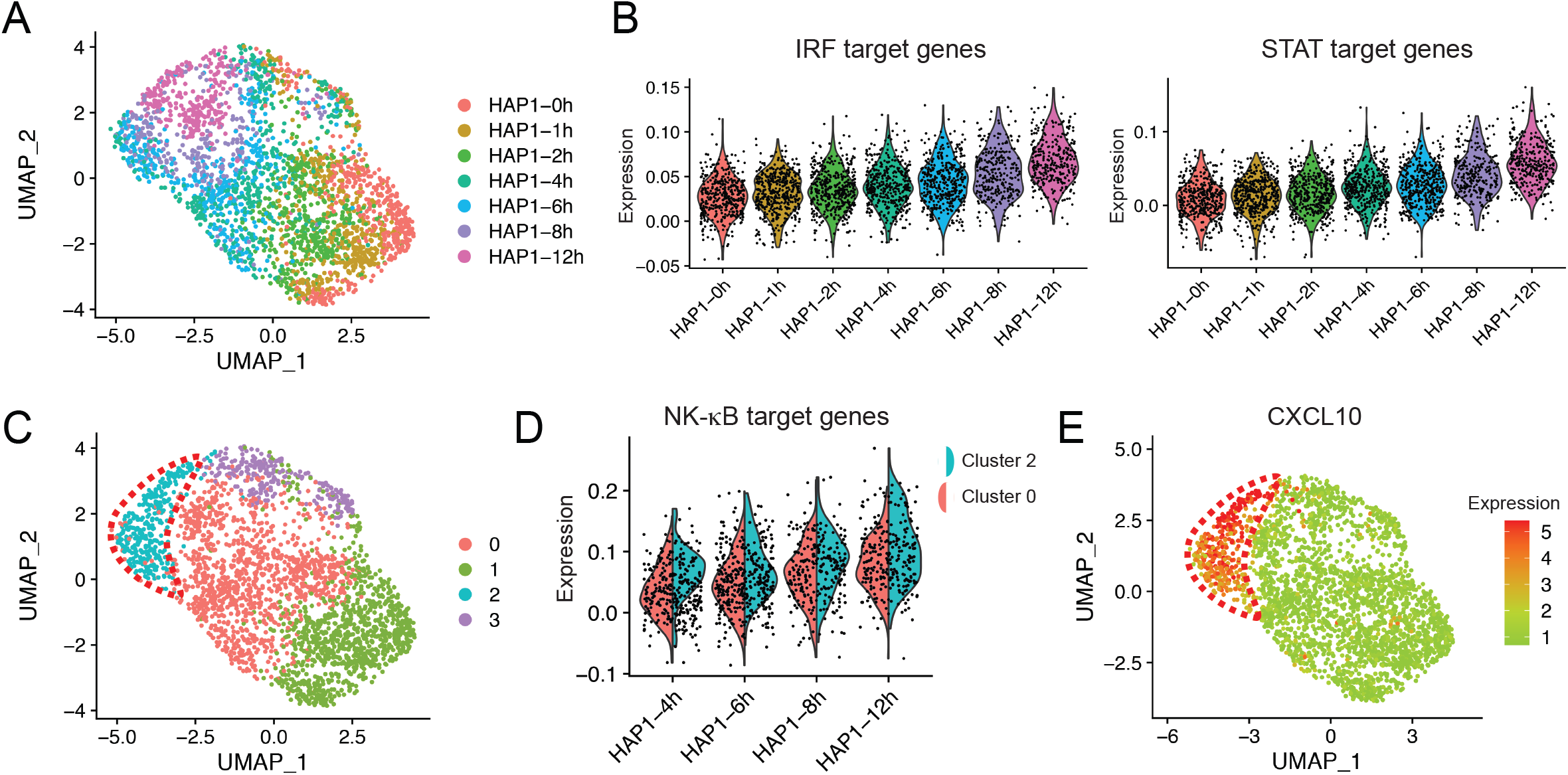
Transcriptomic heterogeneity within HAP1 cells. **A**, Transcriptome-based UMAP of HAP1 cells captured in scRNA-seq, in which cells were colored according to the CASB barcodes. HAP1 cells showed globally a continuous shift from 0 to 12 h. **B**,Violin plots demonstrating the continuous transcriptional activation of predicted IRF and STAT target genes across different cells. **C**,Transcriptome-based UMAP of HAP1, in which cells were unsupervised clustered and colored according to the transcriptomic feature revealed by Louvain algorithm. Cells were clustered into two populations at 4-12 h, one of which exhibited more divergent transcriptome profile from earlier time points. **D**,Violin plots comparing the expression of predicted NF-κB target genes between cluster 0 and 2 at 4-12 h. Predicted target genes were heterogeneously expressed and more actively induced in cluster 2. **E**,Transcriptome-based UMAP of HAP1 cells, in which the expression of CXCL10 were presented with color code and showed activation only in cluster 2 (circled in red) at later time points.

As revealed by unsupervised clustering with Louvain method, cells at later time points (4, 6, 8 and 12 h) were clustered into two populations, one of which exhibited more divergent transcriptome profile from earlier time points (Fig 4C, cluster 2, circled in red). To see whether this is associated with heterogeneous NF-κB activity identified in snATAC-seq, the expression of predicted NF-κB target genes was compared between the two cell populations. Consistent with the heterogeneous NF-κB activity, its target genes also exhibit heterogeneous expression, and were expressed at a higher level in cluster 2 at later time points (Fig 4D). The high induction of CXCL10 and 11, well-known targets of IFN-γ, only in cluster 2 cells further corroborate that the heterogeneous NF-κB activity indeed results in differential responses to IFN-γ in HAP1 cells (Fig 4E&S4C).

## Discussion

CASB is a flexible sample barcoding approach, ready to be prepared in an average molecular biology laboratory. CASB could be used to label cells and nuclei of different cell types and from different species. Moreover, the binding of CASB molecules to the subject is fast and stable, and takes place even at low temperature, which is critical to preserve sample integrity. Importantly, the design of CASB barcoding ssDNA is extremely flexible, which can be easily adapted to different single cell sequencing workflows.

CASB allows scalable sample multiplexing by solely increasing the variety of barcoding ssDNAs. In this study, we tested CASB’s scalability by performing a 20-plex perturbation assay followed by scRNA-seq, which revealed new information about drug response of triple-negative breast cancer cells. Specifically, it demonstrated the different response dynamics of different compounds, and different response of different cell subpopulations to the same drug. The scalability of CASB could be potentially enhanced by combinatorial indexing. When integrated with automated cell handling system, CASB following by scRNA-seq could serve as a powerful platform for single-cell sequencing-based drug screens.

Cell doublets, i.e. two or more cells encapsulated in a same droplet, posed a challenge for single cell sequencing data analysis. Without sample barcoding, cell doublets from the same species could only be estimated mathematically with certain ambiguity (DePasquale, Schnell et al., 2019). To reduce the doublet rate, an often-sought strategy is to limit the number of cells loaded in the microwell- or droplet-based systems. As demonstrated in this study, CASB could reveal cell doublets in high accuracy and as such its application would allow to increase the throughput of single-cell sequencing systems by loading more cells. Indeed, similar strategy has been proposed by using antibody-based barcoding approach (Stoeckius et al., 2018). However, the efficiency of doublet identification correlated to the diversity of sample barcodes. By increasing the sample barcodes to hundreds or even thousands, which could be easily achieved using CASB, we would envisage a much higher doublet detection efficiency, which allows the further optimization of cell loading rate.

Due to the universal presence of glycoprotein on plasma membrane, CASB is applicable to any sample with an accessible plasma membrane. Worth noting, after 1 h transposition reaction at 37 °C, the CASB barcodes remained abundant and showed minimal cross contamination. We believe that CASB can become compatible with many other single-cell sequencing technologies such as CITE-seq (Stoeckius et al., 2017) and SNARE-seq (Chen, Lake et al., 2019), and for samples preserved in different ways such as flash-freezed and formalin-fixed ones. Moreover, by using biotin and fluorophore dual-labeled barcoding ssDNA, one could further enrich cells that are successfully barcoded.

In summary, CASB allows to incorporate additional layers of information into single-cell sequencing experiments. With the ever-increasing throughput of single-cell sequencing technologies, CASB does not only reduce reagent costs, improve data analysis, minimize the batch effect, but also can become a versatile tool in this field by incorporating more diverse types of information, including time-points, treatment conditions and potentially also spatial coordinates. With further improvement, such as using ConA-Streptavidin fusion protein or fluorophore-labeled ssDNA, it will facilitate more novel applications of single cell sequencing technology.

## Materials and Methods

### Experimental materials and methods

#### Cell culture and pre-processing

The MDA-MB-231, MDA-MB-453, T-47D and MCF7 cells were obtained from the ATCC, while HAP1 from Horizon discovery. The MEF cells and mESC were kindly gifted by the Qi Zhou’s and Wei Li’s Lab at the Institute of Zoology, Chinese Academy of Sciences. MDA-MB-231, MDA-MB-453, T-47D, MCF7, HAP1 and MEF cells were cultured in DMEM (11995040, Gibco) with 10% FBS (10270106, Gibco) and 1% P/S (15070063, Gibco) with 5% CO_2_ at 37°C, while the mESC were cultured in Neuralbasal (21103-049, Gibco)-DMEM/F12 (11330-032, Gibco) based medium with N2 (17502048, Gibco), B27(17504-044, Gibco), PD0325901 (s1036, Selleck), Chir99021 (s1263, Selleck) and mLIF (ESG1107, Millipore) with 5% CO_2_ at 37°C. For stimulation with IFN-γ, HAP1 cells were treated with 100 ng/mL IFN-γ (#300-02, Peprotech) for 2, 4, 6, 8 and 12 hours. For scRNA-seq related experiments, cells were tyrpsinized and washed once with PBS, while for snATAC-seq related experiments, after washing with PBS, cells were cryopreserved in 200 μL cryo-medium (10% DMSO, 40% FBS, 50% culture medium) and kept in −80 °C.

#### Compounds and treatment

The compounds used in this study include LCL161 targeting XIAP, Fludarabine inhibiting DNA synthesis, OSI-027 blocking mTOR, Rucaparib and Niraparib targeting PARP1, which were chosen based on their selective inhibitory effect on MDA-MB-231 cells (Garnett, Edelman et al., 2012, Iorio, Knijnenburg et al., 2016). Compounds LCL161, Fludarabine, OSI-027, Niraparib and Rucaparib (HY-15518, HY-B0069, HY-10423, HY-10619 and HY-10617) were obtained from MedChemExpress and dissolved in DMSO. For scRNA-seq experiment, 0.1 μM of LCL161, 0.15 μM of Fludarabine, 2.5 μM of OSI-027, 15 μM of Rucaparib and 12.5 μM of Niraparib were used to treat the cells for 4, 8 or 24 h.

#### Design and synthesis of CASB barcoding ssDNA

For measuring the number of ssDNA molecules immobilized on cell or nuclear membrane, a 5′-biotinylated ssDNA with 5′ and 3′ PCR handles flanking a N8 random sequence was designed: 5′-TCGTCGGCAGCGTCAGATGTGTATA-NNNNNNNN-TATACACATCTCCGAGCCCACGAGAC-3′. For scRNA-seq related experiments, 5′-biotinylated ssDNAs with a 5′ PCR handle followed by a N8 barcode and a 30 nt poly-A tail were designed: 5′-GCTGCGCTCGATGCAAAATA-NNNNNNNN-BAAAAAAAAAAAAAAAAAAAAAAAAAAAAAA-3′. For snATAC-seq related experiments, a 5′-biotinylated forward primer (5′-TCGTCGGCAGCGTCAGATGTGTATAAGAGACAG-CTTGTGGAAAGGACGAAACACCG-3′) and a revers primer (5′-GTCTCGTGGGCTCGGAGATGTGTATAAGAGACAG-GTGTCTCAAGATCTAGTTACGCCAAGC-3′) were used to amplify 222 bp fragments from CROPseq-Guide-Puro plasmids (#86708, Addgene) which have been inserted with different gRNA sequences. To generate ssDNAs, purified PCR products were denatured at 95 °C for 2 mins and immediately put on ice. Information of CASB barcode sequences and their corresponding samples can be found in Table EV2.

#### Assembly of CASB barcode

Biotinylated ConA (C2272, Sigma-Aldrich) and streptavidin (CS10471, Coolaber) were dissolved in 50% glycerol at concentration of 1.6 μM and store in −20 °C, while different biotinylated ssDNAs were diluted at concentration of 100 nM in nuclease-free water and stored in −20 °C. To assemble CASB barcode, streptavidin was firstly mixed with biotinylated ssDNA at molar ratio of 4:1 and incubated for 10 min at room temperature. Afterwards, biotinylated ConA was added to the streptavidin-ssDNA mix at molar ratio of 1:1 and incubated for 10 min at room temperature.

#### Cell labeling with CASB

Half a million cells were collected and suspended in 0.5 mL PBS. Indicated amount of CASB barcode was added to the cells and incubated for 10 mins on ice after thorough mixing. For labeling mCherry+ MEF cells, 2.5 μL assembled CASB was used. For RT-qPCR and scRNA-seq, 5 μL assembled CASB barcode was used. Afterwards, cells were washed once with PBS and then subjected to direct qPCR, direct RT reaction or mixed for scRNA-seq. For qPCR quantification, three independent biological replications were performed for each experiment.

#### Nuclei labeling with CASB

Cells were thawed by adding 800 μL warm culture medium and collected by centrifugation. Afterwards, cells were resuspended in 0.5 mL nuclei extraction buffer (NUC101-1KT, Sigma-Aldrich), incubated for 5 min on ice, and collected by centrifugation (500 g). Extracted nuclei were incubated again with 0.5 mL nuclei extraction buffer, in which indicated amount of CASB barcode was added, for another 5 min on ice. For snATAC-seq, 2.5 μL assembled CASB barcode was used. Nuclei were then washed once with nuclei wash buffer (10 mM Tris 7.4, 10 mM NaCl, 3mM MgCl_2_, 1% BSA, 0.1% Tween-20) and then subjected to direct qPCR or mixed for snATAC-seq. For qPCR quantification, three independent biological replications were performed for each experiment.

#### Quantification of ssDNA immobilized on cell or nuclear membrane

For all quantification experiments using qPCR, standard curves were always first drawn using serially diluted pure ssDNA for calculating the precise number of ssDNA in each reaction. For each reaction of qPCR, 2000 labeled cells in 5 μL PBS were directly mix with 5 μL of primer mix (1 μM) and 10 μL of qPCR master mix (11201ES03, Yeasen). Primers used for quantifying barcoding ssDNA are 5′-TCGTCGGCAGCGTCAG-3′ and 5′-GTCTCGTGGGCTCGGAG-3′.

For measuring the ssDNA with the polyA tail, 20000 cells were directly lysed in 6 μL of 0.17% TrionX-100 (T8787, Sigma-Aldrich) for 3 min at 72 °C, and then revers transcribed with 1^st^ strand cDNA synthesis kit (11119ES60, Yeasen) using RT primer 5′-CACGCACTGACTGACAGAC-TTTTTTTTTTTTTTTTTTTTTTTTTTTTTTV-3′ (final concentration 2.5 μM). The qPCR was performed with ssDNA-specific forward primer 5′-GCTGCGCTCGATGCAAAATA-3′ and Actb-specific forward primer 5′-GTGACAGCATTGCTTCTGTGTAAAT-3′ combining with common reverse primer 5′-CACGCACTGACTGACAGACT-3′.

#### scRNA-seq and snATAC-seq

The scRNA-seq experiments were performed according to the standard protocol of single cell 3′ reagent kits v2 (PN-120237, 10X Genomics) with following modifications. During cDNA amplification, additional primer (5′-GCTGCGCTCGATGCAAAATA-3′, 0.1 μM) was added to amplify CASB barcode. To capture amplified CASB barcode, during “post cDNA amplification reaction cleanup”, the amplified full-length cDNA library was purified with 2X SPRIselect Reagent (B23318, Beckman Coulter) and eluted in 40 μL of nuclease-free water, 10 μL of which was subject to PCR with primer pair 5′- AATGATACGGCGACCACCGAGATCT-ACACTCTTTCCCTACACGACGCTCTTCCGATCT-3′ and 5′-CAAGCAGAAGACGGCATACGAGAT-CTGATC-TGACTGGAGTTCAGACGTGTGCTCTTCCGATCT-GCTGCGCTCGATGCAAAATA-3′ using PrimeSTAR Max PCR master mix (R045A, Takara) to add sequencing adapter sequences to CASB barcode.

The snATAC-seq experiments were performed according to the standard protocol of single cell ATAC reagent kits (PN-1000111, 10X Genomics) with no modification.

#### Next generation sequencing

All sequencing experiments were performed with Illumina NovaSeq 6000 System. The service for scRNA-seq was provided by Haplox genomics center, while for snATAC-seq by Genergy Bio. For scRNA-seq, paired-end 150 bp with i7 8 bp sequencing strategy was used, while for snATAC-seq, paired-end 150 bp with i7 8 bp and i5 16 bp sequencing strategy was applied. All sequencing data were submitted to GEO under the accession number GSE153116.

### Computational methods

#### CASB barcode analysis

For scRNA-seq, raw barcode library FASTQ files were converted to barcode UMI count matrix using custom script leveraging the pysam (Li, Handsaker et al., 2009) package (https://github.com/pysam-developers/pysam). This procedure was similar with a previous method (McGinnis et al., 2019). Briefly, raw FASTQ files were first parsed to use only the reads where the first 16 bases of R1 perfectly match any of the cell barcodes predefined by Cell Ranger. Then, reads where the 20-28 bases of R2 align with at most 1 mismatch to any predefined sample barcodes were used. Reads were grouped by cell barcodes and duplicated UMIs were identified as reads where 17-26 bases of R1 exactly matched.

In snATAC-seq, sample barcodes were in R2 reads and cell barcodes were in R1 reads. Reads with sample and cell barcodes were first extracted from raw FASTQ files of snATAC library using custom script to get the cell-by-sample count matrix.

HTODemux method (Stoeckius et al., 2018) implemented in Seurat package was used to define ‘doublets’, ‘singlets’ and ‘negatives’.

#### scRNA-seq gene expression analysis

FASTQ files were processed using Cell Ranger (10X Genomics, v3.1.0). The reads were aligned to the concatenated hg38-mm10 or hg38 reference using STAR (Dobin, Davis et al., 2013). Cell-associated barcodes were defined by Cell Ranger. Gene expression UMI count matrix (h5 file) was obtained using Cell Ranger with default parameters.

After the pre-processing, RNA UMI count matrices were prepared for scRNA-seq analysis using the ‘Seurat’ R package (Butler, Hoffman et al., 2018). Cells with no more than 4000 reads or 2000 expressed genes were removed. Outlier cells with elevated mitochondrial gene expression were visually defined and discarded. Ribosomal genes and mitochondrial genes were then filtered out.

‘sctransform’ R package (Hafemeister & Satija, 2019) was used to normalize the RNA UMI count data and find highly variable genes. These variable genes were then used during principal component analysis (PCA). Elbow plot was used to select the top principal components. Then these principal components were used for dimensionality reduction with UMAP and unsupervised clustering with Louvain method. Differential gene expression analysis was performed using the ‘FindMarkers’ function in Seurat with ‘MAST’ method (Finak, McDavid et al., 2015). To quantify the magnitude of perturbation induced by drug on gene expression, we compared the proportion of each cell’s k (k=9) nearest neighbors in principal component space with the ‘knn.covertree’ R package. The proportion was normalized by the cell numbers of different groups.

#### snATAC-seq data analysis

After filtering out the reads with sample barcodes, Cellranger-atac (version 1.2.0) was used to process the raw FASTQ files. The reads were aligned to hg38 reference using BWA-MEM (Li & Durbin, 2009). The filtered peak-by-cell matrix (h5 file) obtained after cellranger-atac processing was used in the subsequent analysis. The matrix was first binarized. Cells of low quality (no more than 2000 peaks or more than 500000 peaks, percent of reads in peaks < = 30%, percent of peaks in ENCODE black list >5%) were filtered out. Only cells defined by HTODemux as ‘singlet’ were used for subsequent analysis. ATAC peaks with low coverage (less than 50 cells) or ultra-high coverage (more than 2000 cells) were also removed. The binarized count matrix was normalized by term frequency inverse document frequency (TF-IDF). Latent semantic indexing analysis was performed as applying singular value decomposition (SVD) on the normalized count matrix. Only the 2nd-50th dimensions after the SVD were passed to UMAP for 2D visualization. Motif analysis was performed using chromVAR (Schep, Wu et al., 2017).

The predicted target genes of TF were defined by the nearest genes within 100 kb of the activated ATAC peaks with the TF motif at 12h. The activated ATAC peaks were called by using FindMarkers function with parameter test.use=‘LR’. The average expression levels of these predicted target genes were calculated using ‘AddModuleScore’ function in Seurat package.

#### Gene function enrichment and network analysis using IPA software

The commonly up- or downregulated genes in cluster 1 and 2 compared to cluster 0 of untreated MDA-MB-231 cells were subjected to the Ingenuity Pathway Analysis (IPA) (QIAGEN) (Kramer, Green et al., 2014) to gain insights into the gene functions. The “Diseases & Functions” module under the “Expression Analysis” were used for this purpose. In “Diseases & Functions” module, the analysis was restricted to “Molecular and Cellular Functions”.

## Acknowledgments

### Funding

This work was supported by the Shenzhen Science and Technology Program (Grant No. KQTD20180411143432337, JCYJ20190809154407564 and JCYJ20180504165804015) and the National Natural Science Foundation of China (Grant No. 31970601, 31701237 and 31900431). Computational resource was supported by the Center for Computational Science and Engineering of Southern University of Science and Technology.

### Author contributions

W.C. and L.F. developed the concept of the project. L.F., Q.Z., Y.L. and H.C. designed and performed experiments. G.L. performed bioinformatic analysis. Z.S., W.L. and W.W. assisted in performing experiments. W.C., L.F., G.L. and Y.H. reviewed and discussed results. W.C., L.F. and G.L. wrote the manuscript.

### Competing interests

The authors declare no conflict of interest.

### Data and materials availability

All next-generation sequencing data were submitted to GEO under the accession number GSE153116.

## Supplementary Materials

### Supplementary Figures

**Figure EV1. CASB facilitates scRNA-seq sample multiplexing. A**,The number of ssDNA molecules immobilized on mESC nuclei was quantified used qPCR. The amount of ssDNA immobilized on nuclei increased with the increased usage of ConA-streptavidin-ssDNA complex, and reached at least 120,000 molecules per nucleus. **B**,The poly-A ssDNA molecules immobilized on MEF was detected using RT-qPCR. Both poly-A ssDNA and ActB transcripts can be efficiently capture by RT primer. As expected, barcoding ssDNA can be detected by qPCR even without RT reaction. Error bars represent SD. **C**,Heatmap showing the detected levels of each CASB barcode in individual cells in scRNA-seq. A total of 12068 cells with sufficient reads were captured; 3962 cells that contained at least two major barcodes were assigned as cell doublets; 483 cells were assigned as ‘unlabeled’, as expected due to the inclusion of unlabeled MDA-MB-231 cells. **D**,Boxplot demonstrating the number of UMI derived from both CASB barcode and mRNA transcripts in cell doublets and singlets. Comparing with singlets, more UMI derived from both CASB barcode and mRNA transcripts were detected in doublets. **E**,Scatterplot illustrating the significant positive correlation between the number of detected UMI from CASB and endogenous transcripts among individual cells. “R” means Pearson’s correlation coefficient. **F**,Distribution of normalized CABS UMI counts of singlets and individual cell samples. The CASB barcoding manifested a good uniformity of labeling efficiency (5-95 percentile: 2.1% – 21.8%) (upper panel); comparing with human cell samples, MCF7 cells had slightly lower labeling efficiency (lower panel). **G**,Transcriptome-based UMAP comparing labeled and unlabeled untreated MDA-MB-231 cells, in which two cell populations were intermingled. **H**,Scatterplot demonstrating the well correlated gene expression profiles between labeled and unlabeled untreated MDA-MB-231 cells. “R” means Pearson’s correlation coefficient. **I**,t-SNE projection based on the CASB barcode reads captured in scRNA-seq. Cells were colored according to the CASB barcodes, and doublets were marked in black. All 20 sample barcodes can be detected. **J**,Transcriptome-based UMAP of all MDA-MB-231 cells captured in scRNA-seq. Untreated and 24-h treated cells were highlighted. Cells associated with 24 h-treatment of Niraparib, Rucaparib and OSI-027 could be well distinguished from untreated cells, whereas those with LCL161 and Fludarabine could not.

**Figure EV2. Intrinsic heterogeneity of MDA-MB-231 cells. A**,Transcriptome-based UMAP of untreated and OSI-027-treated MDA-MB-231 cells. Cells were unsupervised clustered into three distinct groups with Louvain method. **B**,Neighbor proportion analysis of untreated and OSI-027-treated MDA-MB-231 cells. In cluster 0, untreated cells were distant from treated cells, while, in cluster 1 and 2, untreated cells were 50% neighbored with treated cells. **C**,Transcriptome-based UMAP of untreated MDA-MB-231 cells. Cells were unsupervised clustered into three distinct groups with Louvain method. **D**,Transcriptome-based UMAP of untreated and Niraparib- and Rucaparib-treated MDA-MB-231 cells. Cells were unsupervised clustered into three distinct groups with Louvain method. **E&F**,Function analysis of genes that were commonly up- or downregulated in insensitive cell populations. **E**,Gene function enrichment revealed that these genes were highly enriched in the cellular compromise and movement pathways. **F**,Genes that were upregulated in insensitive cell populations and predicted to promote cell movement, including VIM, SQSTM1, NPM1 and RACK1.

**Figure EV3. CASB facilitates snATAC-seq sample multiplexing. A**,The number of ATAC-barcode molecules immobilized on mESC nuclei was quantified used qPCR. The amount of ssDNA immobilized on nuclei increased with the increased usage of ConA-streptavidin-ssDNA complex, and could reached at least 80,000 molecules per nucleus. Error bars represent SD. **B**,Heatmap showing the detected levels of each CASB ATAC-barcode in individual cells in snATAC-seq. A total of 2890 cells were obtained with sufficient reads, 305 of which were identified as cell doublets and 23 cells were unlabeled. **C**,ATAC-based UMAP of MDA-MB-231 and HAP1 cells. Cells were colored according to the cell line specific barcodes. MDA-MB-231 cells with its specific CASB barcode presented as an isolated cluster.

**Figure EV4. CASB helps to reveal the dynamic transcriptome change in HAP1 cells. A**,Heatmap showing the detected levels of each CASB barcode in individual cells in scRNA-seq. A total of 3407 cells were captured, 294 of which were identified as cell doublets and 9 cells were unlabeled. **B**,t-SNE projection based on the CASB barcode reads captured in scRNA-seq. Cells were colored according to the CASB barcodes, and doublets were marked in black. **C**,Transcriptome-based UMAP of HAP1 cells, in which the expression of CXCL11 were presented with color code. At later time points, CXCL11 were only actively induced in cluster 2 (circled in red).

### Supplementary Tables

**Table EV1. Genes commonly up- or down-regulated in cluster 1 and 2 of untreated MDA-MB-231 cells.**

**Table EV2. CASB barcode sequences for samples in scRNA/snATAC-seq.**

